# Genome-wide CRISPRi knockdown to map gene essentiality landscape in coliphages λ and P1

**DOI:** 10.1101/2023.05.14.540688

**Authors:** Denish Piya, Nicholas Nolan, Madeline L. Moore, Luis A. Ramirez Hernandez, Brady F. Cress, Ry Young, Adam P. Arkin, Vivek K. Mutalik

## Abstract

Phages are one of the key ecological drivers of microbial community dynamics, function and evolution. Despite their importance in bacterial ecology and evolutionary processes, phage genes are poorly characterized, hampering their usage in a variety of biotechnological applications. Methods to characterize such genes, even those critical to the phage life cycle, are labor-intensive and are generally phage-specific. Here, we develop a systematic gene essentiality mapping method scalable to new phage-host combinations that facilitate the identification of non-essential genes. As proof of concept, we use a catalytically inactive Cas12a mediated genome-wide CRISPRi assay to determine the essential genes in the canonical coliphages λ and P1. Results from a single panel of CRISPRi probes largely recapitulate the essential gene roster determined from decades of genetic analysis for lambda and provide new insights into essential and nonessential loci in P1. We present evidence of how CRISPRi polarity can lead to false positive gene essentiality assignments and recommend caution towards interpreting CRISPRi data on gene essentiality when applied to less studied phages. Finally, we show that we can engineer phages by inserting DNA barcodes into newly identified inessential regions, which will empower processes of identification, quantification and tracking of phages in diverse applications.

## Introduction

Bacteriophages (phages) are the most abundant biological entities on earth and are postulated to play a crucial role in environmental nutrient cycles, agricultural productivity and human health ^1,2^. The full scope of the roles phages play in regulating the activity and adaptation of microbial communities is still emerging^3,4,5^. Phages represent one of the largest pools of genetic diversity with unexplored functional information^6–9^. For example, the majority of phage genes (>70-80%) identified by bioinformatic analysis are of unknown function and show no sequence similarity to characterized genes^10^. Homology-based approaches to connect phage genes to their function are limited by the lack of experimental data ^11,12^. While focused biochemical and genetic analysis are the gold standard for assessment of gene functions, most of these methods are not scalable to the vast amount of new genes being discovered^10^. Unless we develop methods to fill the knowledge gap between phage genetic diversity and gene function, we will be seriously constrained in understanding the mechanistic ecology of phages in diverse microbiomes and harness them as engineerable antimicrobials and microbial community editors^13,14^.

Gaps in phage gene-function knowledge exist even for some of the most well-studied canonical phages^15,16^. Nevertheless, the application of classical phage genetic tools to a few canonical phages over the last few decades has paved the way for generating foundational knowledge of the phage life cycle^15,17,18^. A number of recent technological innovations have also addressed the growing knowledge gap between phage-gene-sequence and the encoded function^19,20,21^. These innovations range from classical recombineering methods ^22,23^ and new phage engineering platforms ^24–28^ to genome editing tools such as CRISPR systems, with or without recombineering technology to create individual phage mutants^13,25,29–33^. Importantly, no method for assessing essentiality without genome modification has been reported. As such, the field is in need of genome-wide technologies that can be used rapidly across diverse phages to assess gene function^14^. At minimum, such a method would provide the foundational knowledge of which phage genes are essential for its infection cycle in a given host, a prerequisite for understanding host range and for engineering.

Catalytically inactive CRISPR RNA (crRNA)-directed CRISPR endonucleases or CRISPR interference (CRISPRi) technology has emerged as a facile tool for carrying out genome-scale targeted interrogation of gene function in prokaryotic and eukaryotic cells without modification of the genome^34,35^. A catalytically inactive or ‘dead’ Cas protein (such as dCas9 or dCas12a) enables programmable transcriptional knock-down (by binding to DNA and forming a transcriptional road block) yielding a loss-of-function phenotype in a DNA sequence-dependent manner^36–40^. Recent work demonstrated that dCas12a is capable of inhibiting infection by phage λ when targeting the essential gene *cro*, suggesting that application of dCas12a with arrayed crRNAs might facilitate genome-wide fitness measurements in phages^41^. The ability to effectively block transcription at target sites distant from promoters makes dCas12a potentially well-suited for repressing transcription of phage genes within operons that show overlapping genetic architecture^15,17,18,42,43^ and those that are highly regulated or vary in expression levels ^44–46^ in a non-competitive plaque assay.

Here, we adopted catalytically inactive Cas12a (dCas12a) to carry out systematic genome-wide interference assays in two canonical phages. The first is coliphage lambda, arguably the best characterized virus in terms of individual gene function and developmental pathways^17^. The second is coliphage P1, which as a powerful generalized transducing phage was instrumental in the development of *E. coli* as a primary genetic model^47^. Its genome is also well annotated but less experimentally characterized than lambda. We first benchmark the CRISPRi technology by applying it to a known set of essential and non-essential genes in both phages, and then extend it genome-wide to query essentiality of all genes in both phages. Although some ambiguities are revealed and significant polarity effects are detected, the method is clearly demonstrated to be applicable to the rapid assignment of non-essential loci in phages, thus paving the way for systematic genome-scale engineering in a variety of applications.

## Results

### Setting up CRISPRi assay targeting phage genes

To ascertain that dCas12a can repress phage gene expression, we designed a phage targeting CRISPRi plasmid system following earlier work ^48^ by expressing both dCas12a and a CRISPR RNA (crRNA) to target specific genes (Methods). Briefly, we placed dCas12a under a anhydrotetracycline (aTc)-inducible Tet promoter and the CRISPR array including the phage targeting crRNA under a strong constitutive promoter on a medium copy plasmid. We then selected a set of known essential and non-essential genes that encode proteins needed at different copy numbers for lambda and P1 (Fig. 1). For lambda, we chose *E*, which encodes the major capsid protein and *Nu1*, which encodes the small terminase subunit. For P1, we chose genes *23, pacA*, and *sit*, encoding the major capsid protein, large terminase subunit, and tape-measure protein, respectively^17,47^. In addition to these essential phage genes, we also chose non-essential P1 genes such as *ppp, upfB* or *ddrB^49^*. We identified Cas12a protospacer adjacent motif (PAM) sites (TTTV) in the 5’ end of the genes (∼20% downstream of the start site) and used 28 bp nucleotide sequence immediately downstream of the PAM site in the coding strand as the spacer region for designing crRNAs.

**Figure 1:**
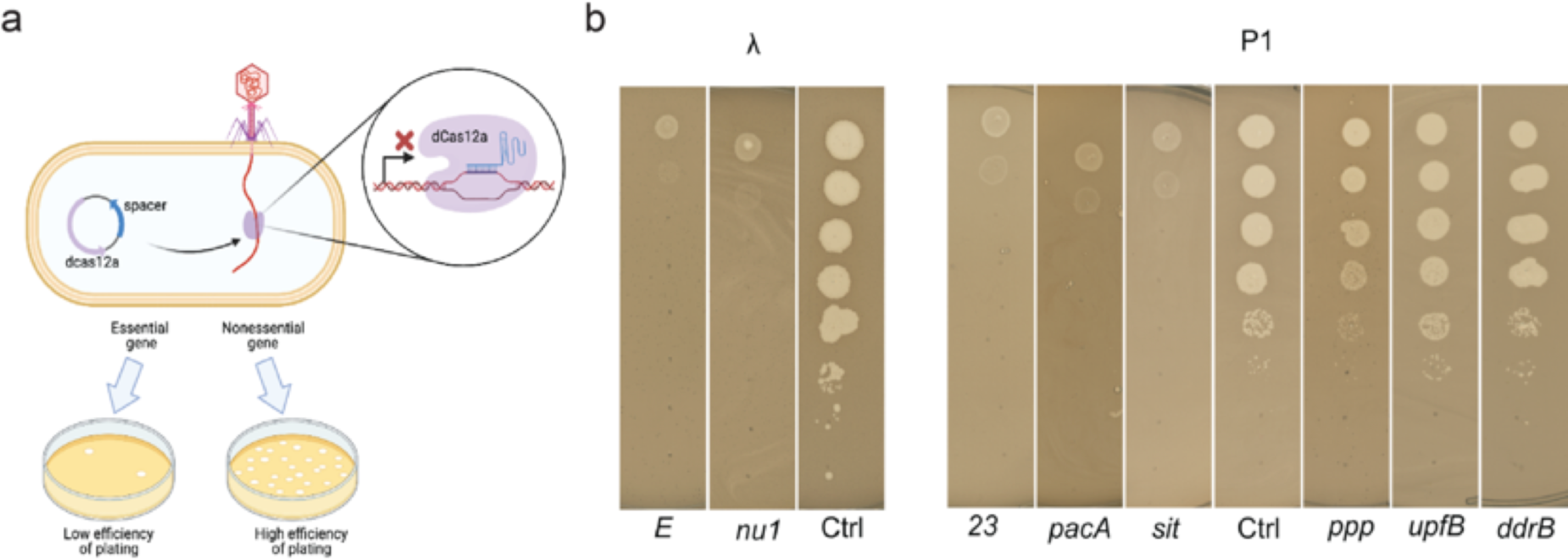
Design and testing of CRISPRi knockdowns to assess gene essentiality in phages lambda and P1. (a) Schematic of CRISPRi assay system (b) Representative images of plaque assays to validate the dCas12a CRISPRi system using gene targets with known essentiality. We employed gRNAs targeting 2 essential genes of phage λ: genes encoding major capsid protein (E) or DNA packaging subunit (Nu1). For phage P1, we used gRNA targeting 3 essential genes: encoding the major capsid protein (gene *23* encoding Mcp), DNA packaging subunit (PacA) and tape measure protein (Sit); and 3 non-essential genes: *ppp, upfB* or *ddrB.* For comparison, phage plaques appearing on an *E. coli* BW25113 lawn expressing a non-targeting crRNA as a control are shown for both phages (Ctrl).

We performed plate-based CRISPRi efficiency assays by moving each variant of the CRISPRi plasmid into *E. coli* BW25113 separately and induced the expression of dCas12a before plating serial dilution of the two phages (Methods). After overnight incubation we compared the plating efficiency on lawns expressing the gene-targeting crRNAs versus a control lawn in which the crRNA did not target either phage (Fig. 1B). We observed that induction of CRISPRi targeting essential genes *E* and *nu1* of lambda and *mcp, pacA* and *sit* of P1 all showed 10^5^ to 10^6^ fold reduction in plating efficiency, whereas targeting nonessential genes *ppp, upfB* or *ddrB* of P1 did not. Overall our CRISPRi benchmarking assays indicated that the dCas12a CRISPRi platform can be used to assess essentiality of phage genes during the infection cycle confirming earlier observations^41^.

### Genome-wide CRISPRi to map gene essentiality in λ

To extend our initial observations to systematically query gene essentiality at genome-wide levels, we considered λ as our pilot case, since it is the most deeply characterized phage with detailed assessments of gene functions well represented in the literature^17,50^. Decades of work on suppressible nonsense mutants of λ phage have helped to define 28 genes (out of total 73 open reading frames (ORF)) as essential for phage growth (Table 1) providing a well-characterized test-bed for validation of our genome-wide CRISPRi assay.

**Table 1:** Gene essentiality mapping of lambda genome. E: essential and NE: nonessential; NT: not tested.

| Locus_tag | Gene | function/Protein name<sup>17</sup> | EOP_average | S.D. | This work | Literature |
| --- | --- | --- | --- | --- | --- | --- |
| lambdap01 | nu1 | DNA packaging protein | 5.5E-06 | 2.1E-06 | E | E<sup>84</sup> |
| lambdap02 | A | TerL | 1.3E-06 | 4.7E-07 | E | E<sup>85,86</sup> |
| lambdap03 | W | gpW family protein | 3.0E-05 | 1.4E-05 | E | E<sup>86,87</sup> |
| lambdap04 | B | portal | 2.5E-05 | 2.1E-05 | E | E<sup>85,86</sup> |
| lambdap05 | C | S49 family peptidase/capsid component; Viral protease | clearing on E-1 |  | E | E<sup>86,88</sup> |
| lambdap06 | nu3 | scaffolding protein | clearing on E-1 |  | E | E<sup>86,89</sup> |
| lambdap07 | D | head decoration protein | 6.8E-06 | 2.4E-07 | E | E<sup>86,90</sup> |
| lambdap08 | E | major capsid protein | clearing on E-1 |  | E | E<sup>86,90</sup> |
| lambdap09 | Fi | DNA packaging protein FI | 2.0E-05 | 0.0E+00 | E | E<sup>86,91</sup> |
| lambdap10 | Fii | head-tail joining protein | clearing on E-1 |  | E | E<sup>87</sup> |
| lambdap11 | Z | tail protein | 2.5E-05 | 7.1E-06 | E | E<sup>50</sup> |
| lambdap12 | U | tail protein | 3.5E-05 | 2.1E-05 | E | E<sup>50</sup> |
| lambdap13 | V | tail protein | 1.7E-05 | 1.9E-05 | E | E<sup>50</sup> |
| lambdap14 | G | minor tail protein G | clearing on E-1 |  | E | E<sup>92</sup> |
| lambdap15 | T | tail assembly protein T | 1.2E-05 | 1.2E-05 | E | E<sup>92</sup> |
| lambdap16 | H | tail tape measure protein | 1.0E-06 | 9.4E-07 | E | E<sup>92</sup> |
| lambdap17 | M | tail protein | clearing on E-1 |  | E | E<sup>92</sup> |
| lambdap18 | L | minor tail protein L | clearing on E-1 |  | E | E<sup>92</sup> |
| lambdap19 | K | tail protein | 8.7E-05 | 1.2E-04 | E | E<sup>92</sup> |
| lambdap20 | I | tail component | 2.6E-05 | 2.7E-05 | E | E<sup>93</sup> |
| lambdap21 | J | host specificity protein J | 1.1E-05 | 1.2E-05 | E | E<sup>85</sup> |
| lambdap26 | lom | Outer membrane beta-barrel protein Lom | 8.8E-01 | 5.3E-01 | NE | NE<sup>17</sup> |
| lambdap27 | stf | protail fiber N-terminal domain containing protein | 3.0E+00 | 2.8E+00 | NE | NE<sup>57</sup> |
| lambdap90 | orf206b | hypothetical protein | 4.3E+00 | 1.1E+00 | NE | NE<sup>57</sup> |
| lambdap28 | tfa | tail fiber protein | 6.3E+00 | 5.3E+00 | NE | NE<sup>57</sup> |
| lambdap29 | orf-194 | tail fiber assembly protein | 3.3E+00 | 1.1E+00 | NE | NE<sup>57</sup> |
| lambdap80 | ea47 |  | 5.5E+00 | 6.4E+00 | NE | NE<sup>17</sup> |
| lambdap81 | ea31 |  | 1.2E+00 | 1.1E+00 | NE | NE<sup>17</sup> |
| lambdap82 | ea59 | ATP-dependent endonuclease | 1.3E+01 | 1.7E+01 | NE | NE<sup>17</sup> |
| lambdap33 | int | tyrosine-type recombinase/integrase | 2.8E+00 | 3.2E+00 | NE | NE<sup>50</sup> |
| lambdap34 | xis | excisionase | 1.6E+00 | 1.1E+00 | NE | NE<sup>50</sup> |
| lambdap35 |  |  |  |  | NT |  |
| lambdap36 | ea8.5 |  | 1.4E+00 | 1.3E+00 | NE | NE<sup>17</sup> |
| lambdap83 | ea22 | ead/Ea22-like family protein | 2.2E+00 | 2.3E+00 | NE | NE<sup>17</sup> |
| lambdap37 | orf61 | hypothetical protein | 9.7E-01 | 8.1E-01 | NE | NE<sup>17</sup> |
| lambdap38 | orf63 | DUF1382 family protein | 1.3E+00 | 3.8E-01 | NE | NE<sup>17</sup> |
| lambdap39 | orf60a | DUF1317 domin-containing protein | 2.4E+00 | 2.0E+00 | NE | NE<sup>17</sup> |
| lambdap41 | exo | YqaJ viral recombinase family protein | 6.9E-01 | 4.4E-01 | NE | NE<sup>17</sup> |
| lambdap84 | bet | recombination protein Bet | 2.1E+00 | 2.5E+00 | NE | NE<sup>17,94</sup> |
| lambdap42 | gam | host-nuclease inhibitor protein Gam | 1.4E+00 | 1.3E+00 | NE | NE<sup>17,94</sup> |
| lambdap85 | kil | host cell division inhibitory peptide Kil | 1.0E+00 | 0.0E+00 | NE | NE<sup>95</sup> |
| lambdap86 | cIII | protease FtsH-inhibitory lysogeny factor CIII | 1.1E+00 | 3.1E-01 | NE | NE<sup>96,97</sup> |
| lambdap45 | ea10 | DUF2528 family protein | 1.7E+00 | 4.7E-01 | NE | NE<sup>17</sup> |
| lambdap46 | ral | Restriction inhibitor protein ral | 1.1E+00 | 3.2E-01 | NE | NE<sup>55,56</sup> |
| lambdap47 | orf28 | hypothetical protein | 1.3E+00 | 4.7E-01 | NE | NE<sup>17</sup> |
| lambdap48 | sieB | Superinfection exclusion protein B | 1.5E+00 | 7.1E-01 | NE | NE<sup>98</sup> |
| lambdap49 | N | antitermination protein N | 1.8E-04 | 1.2E-04 | E | E<sup>99,100</sup> |
| lambdap53 | rexB | exclusion protein | 1.4E+00 | 1.3E+00 | NE | NE<sup>17,98</sup> |
| lambdap87 | rexA | exclusion protein | 3.8E+00 | 4.0E+00 | NE | NE<sup>17,98</sup> |
| lambdap88 | cl | lysogenic repressor | 5.3E+00 | 6.6E+00 | NE | NE<sup>96,101,102</sup> |
| lambdap57 | cro | lytic repressor | clearing on E-1 |  | E | E<sup>103</sup> |
|  | cII |  |  |  | NT | NE<sup>96,101,102</sup> |
| lambdap89 | O | replication protein | clearing on E-1 |  | E | E<sup>104,105</sup> |
| lambdap61 | P | DNA replication protein | clearing on E-1 |  | E | E<sup>104,105</sup> |
| lambdap62 | ren | protein ren | 1.20E-02 | 7.7E-06 | E | NE<sup>106</sup> |
| lambdap63 | NinB | recombination protein NinB | 3.00E-05 | 5.4E-04 | E | NE<sup>17,60</sup> |
| lambdap64 | NinC | phosphoadenosine phosphosulfate reductase family protein | clearing on E-1 |  | E | NE<sup>17,60</sup> |
|  | ninD |  |  |  | NT | NE<sup>17,60</sup> |
|  | ninE |  |  |  | NT | NE<sup>17,60</sup> |
| lambdap67 | NinF |  | 4.00E-05 | 4.0E-06 | E | NE<sup>17,60</sup> |
| lambdap68 | NinG | recombination protein NinG | 8.0E-06 | 1.1E-04 | E | NE |
|  | ninH |  |  |  | NT | NE<sup>17,60,106</sup> |
| lambdap70 | NinI | serine/threonine phosphatase | 4.0E-06 | 5.8E-06 | E | NE<sup>17,60</sup> |
| lambdap71 | Q | antitermination protein | 1.6E-05 | 6.1E-06 | E | E<sup>60</sup> |
| lambdap73 | orf-64 | hypothetical protein | 1.2E-02 | 9.1E-02 | NE |  |
| lambdap74 | S | holin/anti-holin | 1.0E-05 | 4.6E-05 | E | E<sup>107</sup> |
| lambdap92 | S' | holin/anti-holin | 1.0E-05 | 1.7E-04 | E | E<sup>107</sup> |
| lambdap75 | R | endolysin | 6.0E-02 | 5.1E-02 | NE | E<sup>108</sup> |
| lambdap76 | Rz | I-spanin | 5.0E-04 | 1.5E-01 | E | E<sup>109</sup> |
|  | Rz1 |  |  |  | NT | E<sup>109</sup> |
| lambdap77 | bor | serum resistance lipoprotein Bor | 7.9E-01 | 3.0E-01 | NE | NE<sup>17</sup> |
| lambdap78 | - | putative envelope protein | 1.9E+00 | 1.3E+00 | NE | NE<sup>17</sup> |
| lambdap79 | - | hypothetical protein | 3.0E-04 | 1.1E-04 | E | NE<sup>17</sup> |
| **Assay with pQ plasmid** |  |  |  |  |  |  |
| lambdap62 | ren | protein ren | 1.20E-02 | 1.7E+00 | NE | NE<sup>106</sup> |
| lambdap63 | NinB | recombination protein NinB | 3.00E-01 | 9.1E-01 | NE | NE<sup>17,60</sup> |
| lambdap64 | NinC |  | 1.00E+00 | 1.5E+00 | NE | NE<sup>17,60</sup> |
|  | ninD |  |  |  |  | NE<sup>17,60</sup> |
|  | ninE |  |  |  |  | NE<sup>17,60</sup> |
| lambdap67 | NinF |  | 0.03 | 1.4E+00 | NE | NE<sup>17,60</sup> |
| lambdap68 | NinG | recombination protein NinG | 0.2 | 2.1E-01 | NE | NE |
|  | ninH |  |  |  |  | NE<sup>17,60,106</sup> |
| lambdap70 | NinI | serine/threonine phosphatase | 1 | 1.5E+00 | NE | NE<sup>17,60</sup> |

We designed individual crRNAs targeting 67 out of 73 genes of the lambda genome, using the same criteria as used for the pilot studies (by locating PAM sites in the 20-33% of the way through the CDS region of each gene to account for any possible alternative start sites for genes) (Supplementary Fig. 1). The remaining six genes (*cII, ninD, ninE, ninH, Rz1* and lambdap35) were not tested here due to lack of canonical PAM sites. The designed crRNAs were synthesized as separate pairs of oligos and cloned into the CRISPRi plasmid system downstream of a strong constitutive promoter (Methods). Each of these plasmids encoding crRNAs were arranged in an arrayed format and moved into *E. coli* BW25113 as indicator strains for the plate-based CRISPRi assay to measure the EOP (Fig. 2, described above and Methods). The EOP is a quantitative measure of the knockdown for each guide RNA. We assessed the reproducibility of EOP estimations by carrying out biological replicates (total assays >150) and depicted the average EOP of every gene on the lambda map (Fig. 3, (Table 1).

**Figure 2:**
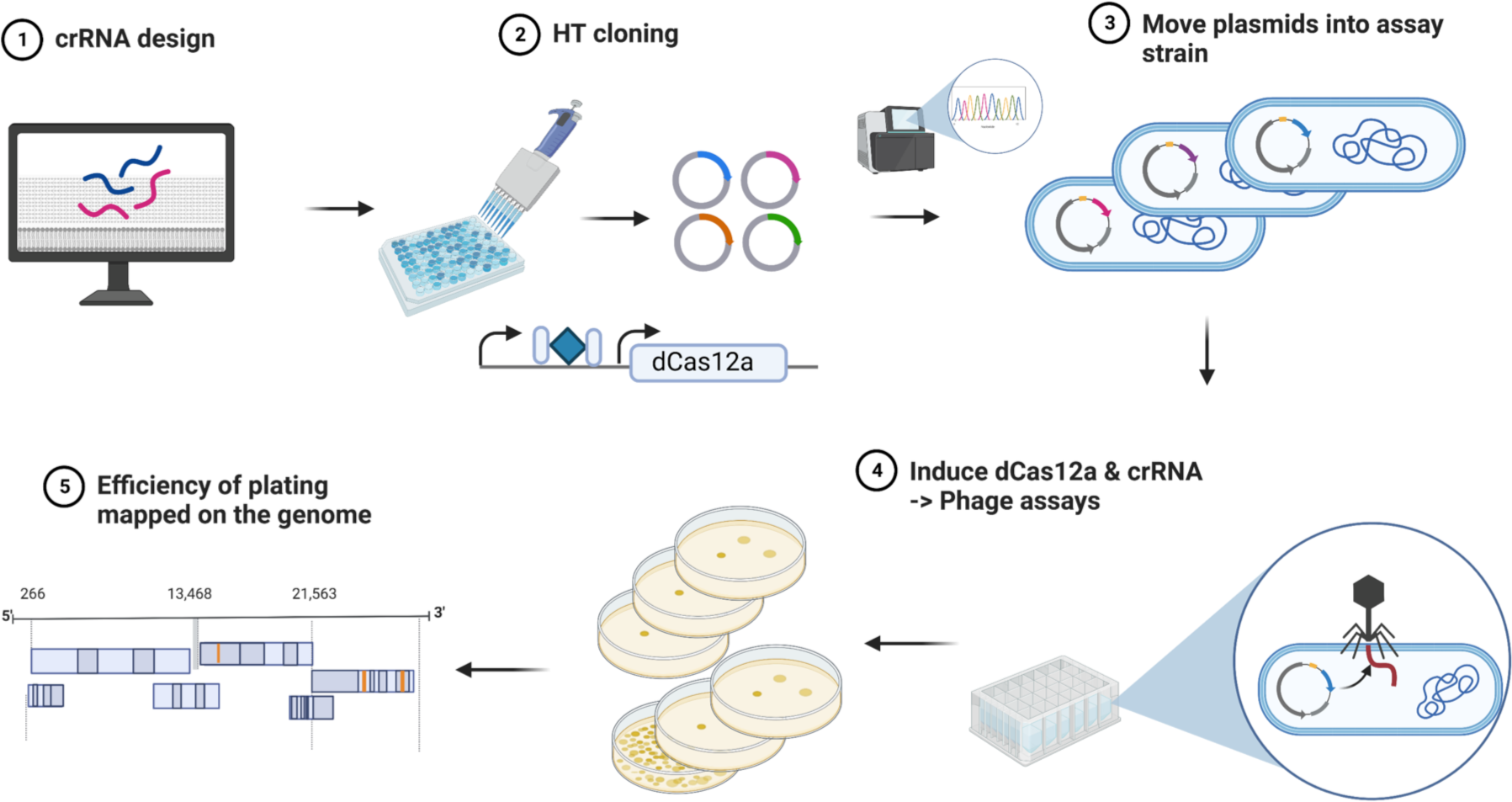
CRISPRi design and assay format: Schematic of steps involved in the arrayed CRISPRi knockdown experiments to assess gene essentiality in phage infectivity cycle.

**Figure 3:**
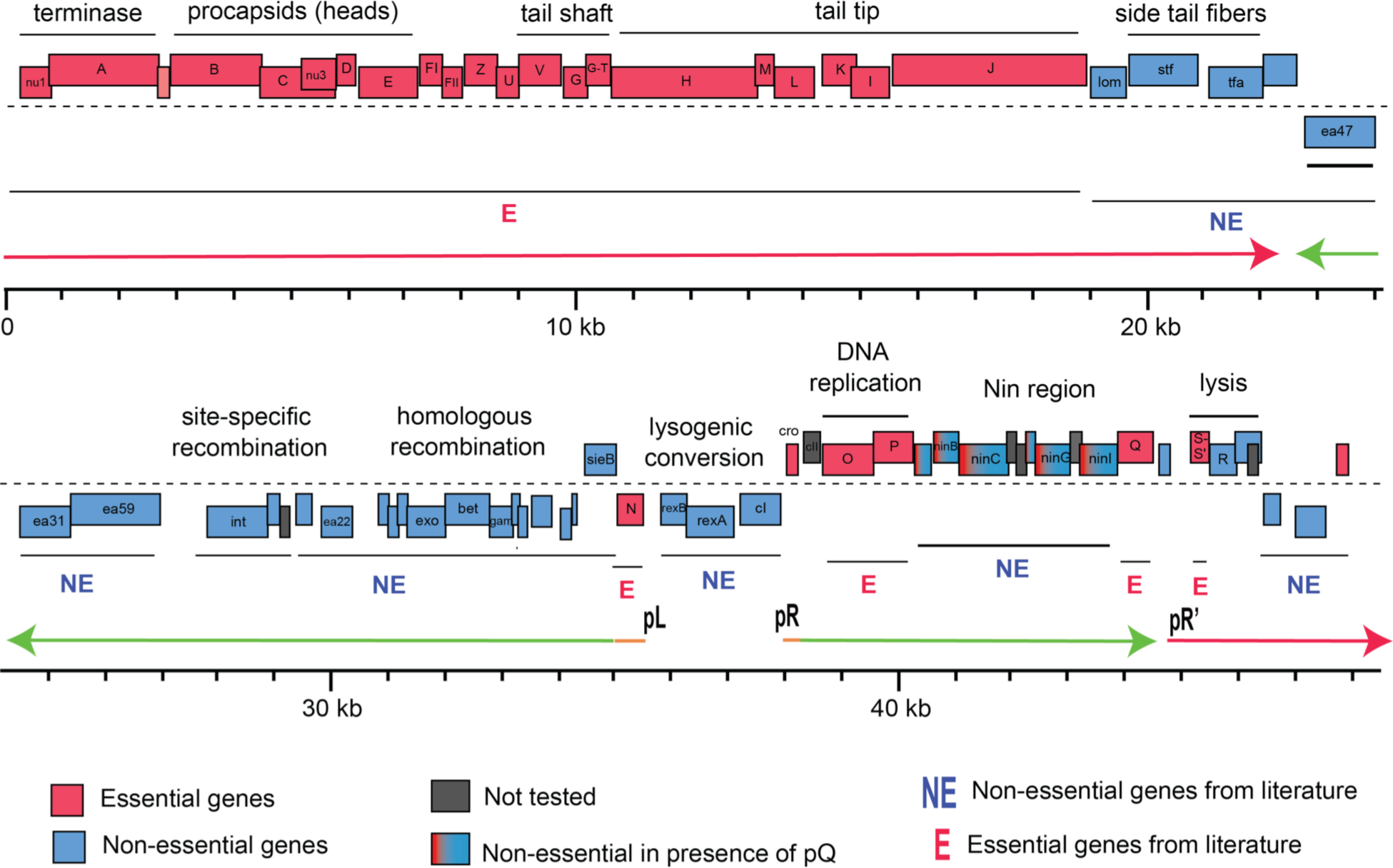
Gene essentiality landscape of phage λ. The genome-wide map of gene essentiality is shown by calculating the EOP as the ratio of plaques appearing on BW25113 lawn expressing crRNA targeting respective lambda phage genes to plaques appearing on BW25113 lawn expressing a nontargeting gRNA. The EOP estimations were done by carrying out biological replicates and depicted the average EOP of every gene on the lambda phage genome map (Methods). Transcripts mentioned in the main text (with promoters) are indicated as thick horizontal arrows: orange, immediate-early transcripts; green, early transcripts; and red, late transcripts.

In total, our CRISPRi assays indicated 35 genes as essential and 32 genes as non-essential. For example, consistent with the literature^17^, knockdown of genes that encode factors involved in the structural assembly of λ virions, either the capsid morphogenesis (Nu1, A, B, C, Nu3, D, E, FI) or tail morphogenesis (V, G, G-T, H, M, L, K, I, J), were detrimental to phage growth with 5-log reduction in EOP. Similarly, repression of genes encoding crucial factors involved in the lytic phase of lambda phage growth cycle, such as transcription antiterminators (proteins N and Q), DNA replication (proteins O and P), transcriptional regulator (Cro) and programmed disruption of host membrane (holin/antiholin S and S’) all showed ∼4-5 log reduction in EOP, indicating their important role in phage fitness phenotype (Table 1).

The longest stretch of dispensable DNA for lambda encompasses > 30% of its genome and is made up of 4 clusters of genes arranged between gene *J* and gene *N* (Fig. 3). These include a cluster of genes *lom-stf-tfa*, 20 genes within pL operon, genes in the immunity region (*rex* and *cI* genes) and genes encoding the lysis program (*R* and *Rz*). We found, except for gene *N*, all genes within pL operon are dispensable for lambda plaque formation (Fig. 3, Table 1). Some of these genes provide functions which would not be expected to have a plaque-formation defect on fully competent lawns, like the superinfection exclusion genes (*rexA, rexB, sieB*)^51^ and genes involved in lysogeny (*int, xis, CIII*)^52^, but others might, such as homologous recombination (*exo, bet, gam*)^53^ and inhibition of host cell division (*kil*)^54^. The knockdown of *ral* (encoding a restriction inhibitor protein) does not result in defect in the EOP because our indicator strain lacks a functional type I restriction system^55,56^. The dispensability of the side tail fiber (which requires *stf* and *tfa*) is in agreement with the known frameshift mutation in the *stf* locus in laboratory strains of λ ^57^.

Interestingly, the CRISPRi-mediated knockdown of a cluster of delayed early genes (*ren, ninB*/*C*/*F*/*G*/*I*) in the pR transcriptional unit indicated that all were essential for plaque-formation, contradicting well-established literature^58–6017^. This ‘*nin* region’ lies between the essential DNA replication genes *O* and *P* and the *Q* gene, encoding the essential late transcription anti-terminator. Phages with a deletion of all the *nin* genes retain full plaque-forming ability^58–60^. The simplest interpretation for this discrepancy is that knockdowns in the *nin* region are polar on transcription of gene Q, the last gene in the transcriptional unit. Polarity has been previously observed for CRISPRi knockdowns in a bacterial context, resulting in false-positives in gene function assignments^61–65^. However, all genes past *cro* are subject to N-mediated anti-termination^17,66^, and to our knowledge, CRISPRi knockdowns have not been tested with phage encoded anti-termination systems. To determine whether the essential phenotype of the *nin* region genes in our assays is due to polarity on gene *Q*, we repeated the knockdown assays on an indicator strain that provides Q in trans from an inducible plasmid^67^. In these conditions, all five genes in the *nin* region targeted by CRISPRi were found to be non-essential, whereas providing Q had no effect on the essentiality of O and P (Fig. 3, Supplementary Fig. 2). We conclude that dCas12a-mediated CRISPRi knockdown repression is insensitive to N-mediated anti-termination. The Q protein is also an anti-terminator and is required for expression of the 27 genes of the late transcript^17,66^. Although most of the genes of this transcript are known to be essential and score that way in our knockdown assays, two of the most promoter-proximal genes score as non-essential, including lambda *orf64* and, to a partial degree, *R*, which shows an intermediate plaque-forming defect. While *R* encodes the endolysin required for lysis, it is known to be produced in great excess, so a significant knockdown might still generate enough bacteriolytic activity to account for the intermediate plaque defect. *Orf64* is indicated to be non-essential^17^, but it is unclear why the knockdown is not polar on the many essential genes downstream.

### Extending genome-wide CRISPRi assay to coliphage P1

We next extended the genome-wide CRISPRi knockdown assays to assess gene essentiality in coliphage P1. The 93 Kbp genome of P1 is composed of 117 genes, organized into 45 transcriptional units, with 8 involved in the lysis-lysogeny switch and plasmid prophage maintenance, while 37 are involved in lytic development^47^. Despite its paradigm status, a large proportion of gene function assignments still awaits experimental verification^47,68^. Early gene expression and the lytic-lysogenic decision are controlled by the primary phage repressor C1 while Lpa (Late Promoter Activator) positively regulates late transcription. There are 11 late promoters, all of which have a conserved 9 bp inverted repeat that serves as the Lpa binding site. Among the 117 genes, 30 have been identified as essential for plaque-formation by amber mutant and targeted deletion methods (Table 2). Experimental evidence for non-essentiality was available for 55 other genes, which makes P1 nearly as good for benchmarking the CRISPR knockdown strategy as lambda.

**Table 2:** Gene essentiality mapping of Phage P1 genome. E: essential and NE: nonessential; NT: not tested; Red genes are structural & essentials. Pink ones are required for marginal essential.

| locus_tag | gene | function<sup>47</sup> | P_average | SD | This work | Literature |
| --- | --- | --- | --- | --- | --- | --- |
| P1_gp002 | cra | cre associated function | 6.6E-01 | 1.3E-01 | NE | NE<sup>110,111</sup> |
| P1_gp003 | cre | cyclization recombinase | 1.2E+00 | 8.1E-01 | NE | NE<sup>110,111</sup> |
| P1_gp004 | c8 | establishment of lysogeny | 1.5E+00 | 3.9E-01 | NE |  |
| P1_gp005 | ref | recombination enhancement | 1.3E+00 | 3.5E-01 | NE | NE<sup>111</sup> |
| P1_gp006 | mat | maturation control | 2.8E-01 | 3.5E-03 | NE | E<sup>112</sup> |
| P1_gp007 | res | restriction component | 1.2E+00 | 3.5E-02 | NE | NE<sup>113</sup> |
| P1_gp008 | mod | modification component | 1.0E+00 | 3.2E-01 | NE | NE<sup>113</sup> |
| P1_gp009 | lxc | modulator of C1 action; | 7.0E-01 | 4.2E-01 | NE | NE<sup>114</sup> |
| P1_gp010 | ulx | enhances incorporation of darB | 6.4E-01 | 1.6E-01 | NE | NE<sup>115</sup> |
| P1_gp011 | darB | antirestriction | 5.8E-01 | 2.5E-01 | NE | NE<sup>115</sup> |
| P1_gp012 | prt | portal | essential |  | E | E<sup>47</sup> |
| P1_gp013 | pro | head processing | essential |  | E | E<sup>47</sup> |
| P1_gp115 | lydE | putative antiholin | 9.8E-01 | 8.8E-01 | NE |  |
| P1_gp014 | lydD | putative holin | 1.1E+00 | 1.0E-01 | NE |  |
| P1_gp015 | lyz | lysozyme | 5.0E-02 | 3.5E-03 | E | E<sup>116</sup> |
| P1_gp016 | ssb | single stranded DNA binding protein | 5.7E-01 | 2.0E-02 | NE |  |
| P1_gp017 | isaA | IS1 insertion-associated gene | 8.6E-01 | 4.7E-01 | NE |  |
| P1_gp018 | insB | IS1 transposition protein | 5.1E-01 | 1.6E-01 | NE |  |
| P1_gp019 | insA | IS1 transposition protein | 1.0E+00 | 2.2E-01 | NE |  |
| P1_gp020 | isaB | IS1 insertion-associated gene | 1.4E+00 | 2.2E-01 | NE |  |
| P1_gp021 | hxr | possible repressor; homolog of Xre | 1.6E+00 | 1.2E+00 | NE | NE<sup>115</sup> |
| P1_gp022 | ddrB | antirestriction | 8.5E-01 | 1.3E-01 | NE | NE<sup>115</sup> |
| P1_gp116 | iddB | internal to ddrB | 5.4E-01 | 2.6E-01 | NE | NE<sup>115</sup> |
| P1_gp023 | ddrA | antirestriction | 9.8E-01 | 3.1E-01 | NE | NE<sup>115</sup> |
| P1_gp024 | darA | antirestriction | 6.5E-01 | 2.9E-01 | NE | NE<sup>115</sup> |
| P1_gp025 | hdf | antirestriction | 1.2E+00 | 8.4E-01 | NE | NE<sup>115</sup> |
| P1_gp026 | lydB | lysis determinant; prevents premature lysis | 2.3E-02 | 2.6E-02 | E | NE<sup>116,117</sup> |
| P1_gp027 | lydA | holin | 1.2E-01 | 2.4E-03 | NE | NE<sup>116,117</sup> |
| P1_gp028 | lydC | holin | 1.1E+00 | 1.2E-01 | NE |  |
| P1_gp029 | cin | site-specific recombinase | 1.4E+00 | 9.5E-01 | NE | NE<sup>118</sup> |
| P1_gp001 | v prime | C-terminal moiety of tail fiber gpS | 5.3E-01 | 3.2E-01 | NE | NE<sup>119</sup> |
| P1_gp030 | J prime | structural protein gpU prime of tail fiber | 8.6E-01 | 8.0E-01 | NE | NE<sup>119</sup> |
| P1_gp031 | U | tail fiber structure or assembly | essential |  | E |  |
| P1_gp032 | S | tail fiber structure or assembly | essential |  | E |  |
| P1_gp033 | R | tail fiber structure or assembly | essential |  | E | E<sup>120</sup> |
| P1_gp034 | 16 | baseplate or tail tube | essential |  | E | E<sup>121</sup> |
| P1_gp035 | bplA | putative baseplate structure, may correspond to gene 3 | essential |  | E |  |
| P1_gp036 | pmgA | Putative morphogenetic function | essential |  | E | E<sup>68</sup> |
| P1_gp037 | sit | putative tape measure protein | essential |  | E |  |
| P1_gp038 | pmgB | Putative morphogenetic function | essential |  | E | E<sup>68</sup> |
| P1_gp039 | tub | tail tube | essential |  | E |  |
| P1_gp040 | pmgC | Putative morphogenetic function | essential |  | E | E<sup>68</sup> |
| P1_gp041 | simC | superimmunity | 1.7E+00 | 6.9E-01 | NE | NE<sup>122</sup> |
| P1_gp042 | simB | superimmunity | 3.8E+00 | 8.4E-01 | NE | NE<sup>122</sup> |
| P1_gp043 | simA | superimmunity | 9.0E-01 | 4.9E-01 | NE | NE<sup>122</sup> |
| P1_gp044 | 4 RNA | acts on icd and ant mRNA | 8.3E-01 | 9.4E-01 | NE |  |
| P1_gp045 | icd | reversible inhibition of cell division | 6.0E-01 | 2.2E-01 | NE | NE<sup>123</sup> |
| P1_gp046 | ant1 | antagonizes C1 repression | 2.0E+00 | 1.1E+00 | NE | NE<sup>124</sup> |
| P1_gp047 | ant2 | product antagonizes C1 | 2.3E+00 | 1.8E+00 | NE | NE<sup>124</sup> |
| P1_gp048 | ask | regulatory region of kilA gene; | 1.2E+00 | 4.9E-01 | NE |  |
| P1_gp049 | kilA | product can kill host | 8.6E-01 | 8.0E-01 | NE | NE<sup>124</sup> |
| P1_gp050 | repL | initiates replication at oriL | 7.1E-01 | 3.0E-01 | NE | NE<sup>124</sup> |
| P1_gp051 | rlfA | possibly associated with lytic replication | 1.3E+00 | 6.1E-02 | NE |  |
| P1_gp052 | rlfB | possibly associated with lytic replication | 8.8E-01 | 3.0E-02 | NE |  |
| P1_gp053 | pmgF | putative morphogenetic function | 1.2E+00 | 1.0E-02 | NE | NE<sup>68</sup> |
| P1_gp054 | bplB | baseplate structure | essential |  | E |  |
| P1_gp055 | pmgG | putative morphogenetic function | essential |  | E | E<sup>68</sup> |
| P1_gp056 | 21 | baseplate or tail tube | essential |  | E | amber mutant reported<sup>121</sup> |
| P1_gp057 | 22 | tail sheath | essential |  | E | amber mutant reported<sup>121</sup> |
| P1_gp058 | 23 | Major head protein | essential |  | E | amber mutant reported<sup>121</sup> |
| P1_gp059 | parB | active partitioning of P1 plasmid during cell division | 1.3E+00 | 7.6E-01 | NE |  |
| P1_gp060 | parA | active partitioning of P1 plasmid during cell division | 1.6E+00 | 1.5E+00 | NE |  |
| P1_gp061 | repA | initiates replication from oriR; plasmid replication | 9.5E-01 | 6.4E-02 | NE |  |
| P1_gp062 | upfA |  | 1.2E+00 | 8.9E-01 | NE | NE<sup>68</sup> |
| P1_gp063 | mlp | membrane lipoprotein precursor | 2.4E-01 | 1.9E-01 | NE |  |
| P1_gp064 | ppfA | possible periplasmic function | 1.3E+00 | 1.2E+00 | NE |  |
| P1_gp065 | upfB |  | 1.3E+00 | 1.0E+00 | NE | NE<sup>68</sup> |
| P1_gp066 | upfC |  | 1.1E+00 | 6.8E-01 | NE | NE<sup>68</sup> |
| P1_gp067 | uhr |  | 8.6E-01 | 1.9E-01 | NE | NE<sup>68</sup> |
| P1_gp068 | hrdC | hypothetical recombination associated protein of RdgC family | 8.7E-01 | 4.6E-01 | NE |  |
| P1_gp069 | dmt-B | DNA methyltransferases; methylates A at GATC | 1.4E+00 | 1.2E+00 | NE | NE<sup>125</sup> |
| P1_gp070 | dmt-A |  | 1.4E+00 | 7.6E-03 | NE |  |
| P1_gt071 | trnT |  | 9.2E-01 | 4.0E-01 | NE |  |
| P1_gp072 | plp | putative lipoprotein | 1.6E+00 | 8.2E-01 | NE |  |
| P1_gp073 | upl |  | 1.9E+00 | 5.9E-01 | NE | NE<sup>68</sup> |
| P1_gp074 | tciA | tellurite or colicin resistance or inhibition of cell division | 6.5E-01 | 6.5E-02 | NE |  |
| P1_gp075 | tciB | tellurite or colicin resistance or inhibition of cell division | 9.9E-01 | 1.0E-01 | NE |  |
| P1_gp076 | tciC | tellurite or colicin resistance or inhibition of cell division | 1.3E+00 | 4.4E-01 | NE |  |
| P1_gt117 | trnI |  | 1.4E+00 | 7.9E-01 | NE |  |
| P1_gp077 | ban | dnaB homolog | 1.2E+00 | 4.8E-01 | NE | NE<sup>126</sup> |
| P1_gp078 | dbn | downstream of ban | 1.3E+00 | 1.3E-01 | NE | NE<sup>68</sup> |
| P1_gp079 | 5 | baseplate | essential |  | E | amber mutant in gene 5. particles with contracted tails<sup>121</sup> |
| P1_gp080 | 6 | tail length | 8.3E-03 | 1.3E-03 | E | amber mutant in gene 6. particles with shorter tails<sup>121</sup> |
| P1_gp081 | 24 | baseplate or tail stability | 4.9E-03 | 5.0E-03 | E | amber mutant in gene 5. polysheath structure present<sup>121</sup> |
| P1_gp082 | 7 | tail stability | essential |  | E | amber mutant in gene 7: several abnormal tail structures present<sup>121</sup> |
| P1_gp083 | 25 | tail stability | 1.1E-01 | 6.6E-02 | NE | amber mutant in gene 25 phenotype similar to gene 7<sup>121</sup> |
| P1_gp084 | 26 | baseplate; | 1.2E-01 | 1.5E-01 | NE | amber mutant phenotype similar to gene 5:<sup>121</sup> |
| P1_gp085 | pmgL | putative morphogenetic function | 1.3E+00 | 9.4E-01 | NE | NE<sup>68</sup> |
| P1_gp086 | pmgM | putative morphogenetic function | 3.8E-01 | 2.1E-01 | NE | NE<sup>68</sup> |
| P1_gp087 | pmgN | putative morphogenetic function | 1.4E-02 | 1.5E-03 | E | NE<sup>68</sup> |
| P1_gp088 | pmgO | putative morphogenetic function | 1.1E+00 | 8.7E-01 | NE | NE<sup>68</sup> |
| P1_gp089 | pmgP | putative morphogenetic function | 7.2E-01 | 7.3E-02 | NE | NE<sup>68</sup> |
| P1_gp090 | ppp | protein phosphatase | 1.2E+00 | 8.0E-02 | NE | NE<sup>68</sup> |
| P1_gp091 | pmgQ | putative morphogenetic function | 1.4E+00 | 7.5E-01 | NE | NE<sup>68</sup> |
| P1_gp092 | pmgR | putative morphogenetic function | 9.4E-01 | 3.0E-01 | NE | E<sup>68</sup> |
| P1_gp093 | pmgS | putative morphogenetic function | 1.2E+00 | 3.2E-01 | NE | NE<sup>68</sup> |
| P1_gp094 | pap | acid phosphatase | 7.7E-01 | 2.3E-01 | NE | NE<sup>68</sup> |
| P1_gp095 | pmgT | putative morphogenetic function | 9.8E-01 | 9.7E-01 | NE | NE<sup>68</sup> |
| P1_gp096 | pmgU | putative morphogenetic function | 2.1E-01 | 6.4E-02 | NE | NE<sup>68</sup> |
| P1_gp097 | pmgV | putative morphogenetic function | 1.9E+00 | 2.0E+00 | NE | NE<sup>68</sup> |
| P1_gp098 | upfM | unknown protein function |  |  | NT | NE<sup>68</sup> |
| P1_gp099 | upfN | unknown protein function | 8.3E-01 | 2.4E-01 | NE | NE<sup>68</sup> |
| P1_gp100 | upfO | unknown protein function | 9.5E-01 | 6.4E-01 | NE | NE<sup>68</sup> |
| P1_gp101 | hot | DNA replication | 1.2E+00 | 1.6E+00 | NE |  |
| P1_gp102 | lrx | LexA-regulated functions | 7.9E-01 | 7.7E-01 | NE |  |
| P1_gp103 | humD | DNA repair | 5.4E-01 | 3.3E-01 | NE |  |
| P1_gp104 | phd | i-toxin of P1 toxin-antitoxin system | 5.5E-01 | 7.1E-02 | NE |  |
| P1_gp105 | doc | toxin of P1 toxin-antitoxin system | 3.6E-01 | 1.2E-02 | NE |  |
| P1_gp106 | pdcA | unknown protein function |  |  | NT | NE<sup>68</sup> |
| P1_gp107 | pdcB | unknown function | 6.8E-01 | 2.6E-01 | NE | NE<sup>68</sup> |
| P1_gp108 | lpa | late promoter activator | essential |  | E | amber mutant described<sup>127</sup> |
| P1_gp109 | pacA | DNA packaging | essential |  | E | amber mutant defective in pac cleavage<sup>128</sup> |
| P1_gp110 | pacB | DNA packaging | essential |  | E | amber mutant defective in pac cleavage<sup>128</sup> |
| P1_gp111 | c1 | lytic repressor | 1.4E+00 | 1.3E+00 | NE |  |
| P1_gp112 | coi | C1 inactivator | 7.4E-01 | 8.4E-01 | NE | NE<sup>129</sup> |
| P1_gp113 | imcB | immunity function | 8.2E-01 | 7.3E-01 | NE |  |
| P1_gp114 | imcA | immunity function |  |  | NT |  |

We designed individual crRNAs targeting 114 out of the 117 genes; the remaining 3 genes (*upfM, pdcA and imcA*) were not tested due to lack of PAM sites. Using the same workflow described for lambda, we found 87 genes as non-essential and 27 genes as essential. (Supplementary Fig. 3, Fig. 3). Five known essential genes were missed by the knockdown screen: *mat, repL, 25, 26*, and *pmgR*. In addition, one gene, pmgN, was found to be essential, in contradiction with the recent deletion analysis survey^68^. From the perspective of identifying non-essential genes, 54 of the 55 genes for which there was some evidence of non-essential character were confirmed by the knockdown. In addition, the knockdown approach demonstrates non-essentiality for a further 33 genes. Taken together, four large segments comprising nearly 60 kb of the P1 genome are occupied by genes dispensable for lytic growth and thus available for specific engineering (Table 2).

### Downstream application of gene essentiality mapping

To demonstrate one downstream application of the knockdown approach to gene-essentiality mapping, we sought to insert an unique DNA tag into both λ and P1 at a gene locus that we found to be dispensable. As DNA barcodes are heritable they can be used for rapid identification of different phage samples by standardizing the workflow, assuming their insertion does not impact phage fitness. Such unique barcoding of different phages could enable quantitative tracking and measure of individual phage fitness in multi-phage formulations in different applications. As a proof of concept, we inserted a unique DNA barcode in genes res (between *cra*-*darB*) and red, of P1 and lambda respectively. With these two *b*c (barcoded) constructs, we tested whether we could quantify different phage combinations. To do this, we mixed barcoded (bc) phage P1-bc and λ-bc in different ratios and subjected them to Barseq PCR sequencing^69,70^. Our Barseq quantification method not only successfully quantified different ratios of barcoded phage P1 and lambda, but also captured the differences in plaque-forming units/ml to barcode abundance (Fig. 5).

**Figure 4:**
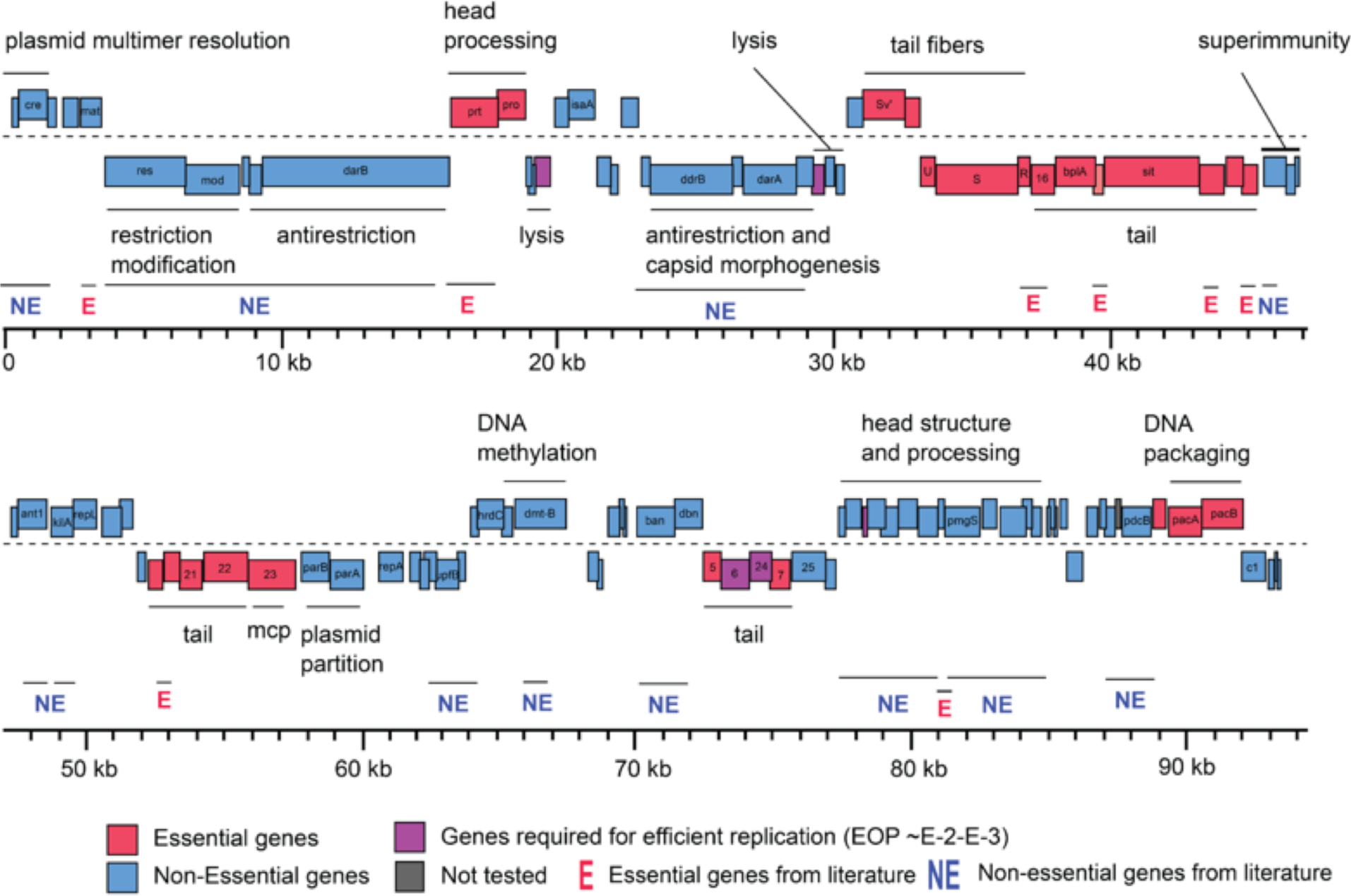
Gene essentiality landscape of phage P1. The genome-wide map of gene essentiality is shown by calculating the EOP as the ratio of plaques appearing on BW25113 lawn expressing crRNA targeting respective P1 phage genes to plaques appearing on BW25113 lawn expressing a nontargeting gRNA. The EOP estimations were done by carrying out biological replicates and depicted the average EOP of every gene on the P1 phage genome map (Methods).

**Figure 5:**
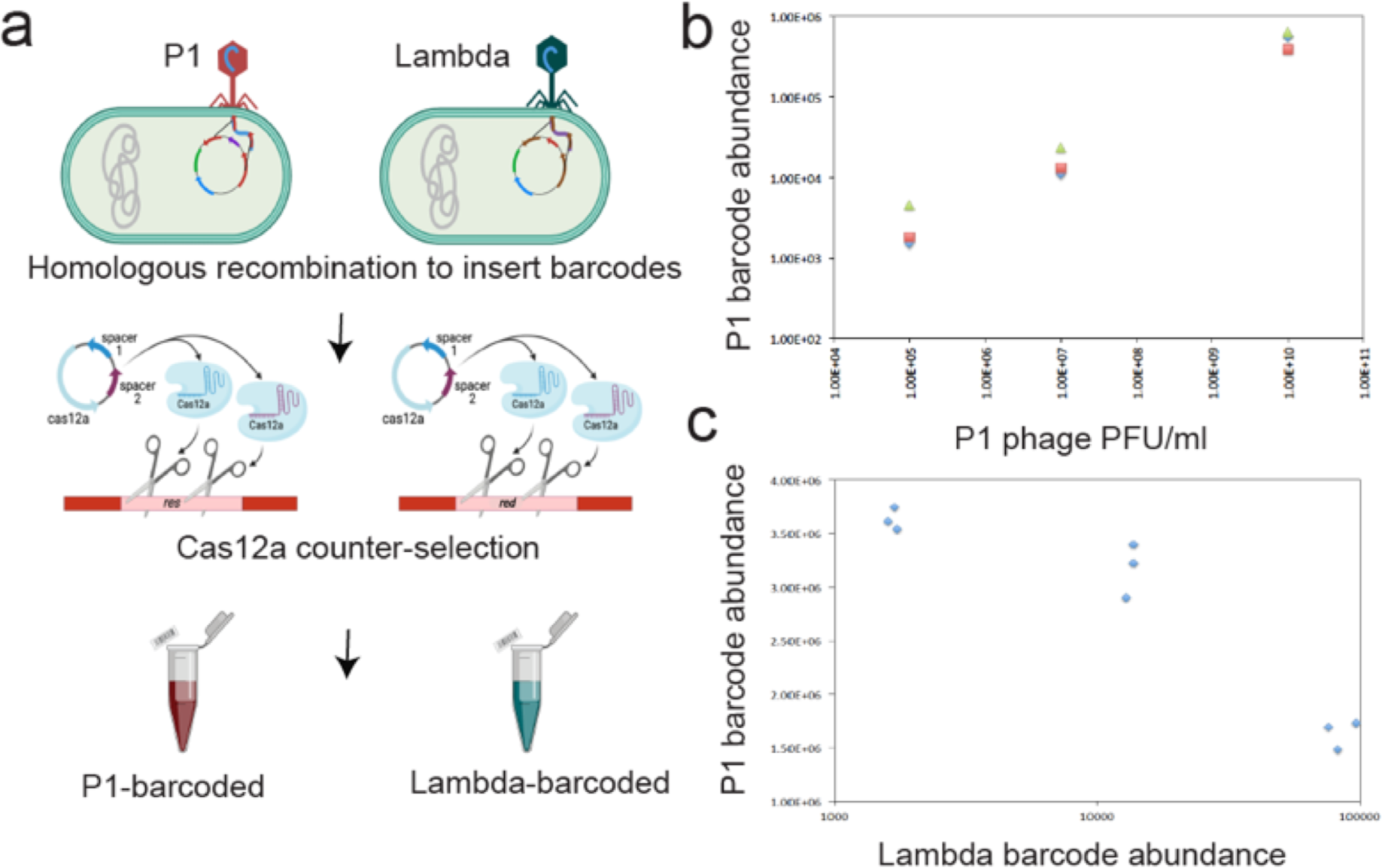
Insertion and quantification of random DNA barcodes on a non-essential genomic location of Lambda and P1vir phage. (a) Schematic of phage engineering approach: homologous recombination method was used to engineer phages with random barcodes at a non-essential genomic loci and nuclease active Cas12a-based counter-selection was used to enrich engineered phages. Schematic is shown for barcode insertion and counter-selection for lambda phage at the *red* locus and P1 phage at *res* locus (b) Barcode abundance of P1 phage against its PFU/ml estimations in triplicates (c) Barcode abundance for both barcoded lambda and P1 phages, when mixed at different ratios. Estimations done in triplicates in a pool (Methods).

## Discussion

CRISPR-based technologies have revolutionized the functional genomics field^34^. CRISPRi in particular has emerged as a major technology for genome-wide mapping of essential and non-essential genes in bacteria^34,35^. Here we assessed the feasibility of using dCas12a system for performing a genome-wide survey of two paradigm phages, lambda and P1, using crRNAs designed to achieve gene-specific “knockdown”. Results from our arrayed CRISPRi assays are consistent with known assignments of gene essentiality in both phages, provide novel insights and present a genome-wide landscape of gene essentiality for phage P1 for the first time to the best of our knowledge (Table 1 and 2, Fig. 3, Fig. 4). Lambda and P1 phages have quite distinct transcriptional organization, making CRISPRi differentially suited for probing stretches of non-essential genes in these phages (below). With an organized map of gene essentiality in hand it is now possible to identify locations in these phage genomes wherein insertion of an exogenous ‘payload’ are less likely to disrupt critical function, as well as longer regions that can be deleted or replaced with custom DNA. We demonstrate this by inserting a DNA barcode into the lambda and P1 genome at discovered inessential loci and confer the ability to track and quantify distinct phages in a mixed phage formulation. Finally, this study uncovers the polar effect of CRISPRi in phages. We recommend using CRISPRi for mapping non-essential regions while caution towards interpreting essential gene assignments when applied to less studied phages where transcripts have not yet been mapped. We discuss these insights below.

Overall, the genome-wide CRISPRi assay results demonstrated dCas12a was effective; that is, nearly every non-essential and essential lambda gene knockdown was scored correctly, and essential lambda genes were scored as essential, based on reduction of plating efficiency by three powers of ten or more in the presence of dCas12a and the cognate crRNA (Fig. 3, Table 1). However, a cluster of delayed early genes in the *nin* region of the pR transcript of lambda were scored as essential despite unambiguous evidence that this entire region can be deleted without impairing the plaque-forming ability of the phage^58–60^. Because of its DNA-binding function, the bound dCas12a/crRNA complex is necessarily a road-block that would be polar on all downstream genes, as confirmed experimentally for the knockdowns of *lacZ* in the *lacZYA* operon of *E. coli^38^*. The reason for this polarity is because this cluster of non-essential *nin* genes is upstream of gene *Q*, which encodes the late-gene activator required for late-gene expression. Thus roadblocks in the *nin* genes should be polar on gene Q. Accordingly, when we supplied Q in trans, the *nin* genes all scored properly as non-essential (Fig. 3). Unfortunately, the same rationale applies to the other genes served by pR (Fig. 3). Thus knockdowns in *cro, O*, and *P* are also polar on *Q*. With *Q* added in trans, all three upstream genes read out as essential but only the gene *P* result is confirming, since the *cro* and *O* knockdowns should be polar on *P*. The situation is better for the pL transcriptional unit because the only essential gene is the first one, *N*. Thus for pL, all 19 genes that were tested are scored correctly, as non-essential.

Similar challenges for CRISPRi essentiality determination are noted for the late genes, expressed from pR’ in a 27kb mRNA (Fig. 3). Twenty-one genes from *Nu1* through *J* read out correctly as essential, but since the first twenty knockdowns should be polar on J, nothing can be concluded for their essentiality. Moreover, the results for the upstream genes *orf64* and *R* are confounding. From the same perspective as used on the pR transcript, knockdown roadblocks in all the upstream genes in this transcriptional unit should be read out as essential. This was observed for gene *p79* (which is non-essential, but essential in our assay), but it was not observed for the knockdowns of *orf64* and *R*. In CRISPRi studies on bacterial genomes^61–65,71^ similar polarity issues have been noted, and contradictions have been explained by invoking the presence of cryptic promoters downstream of the roadblock site^64^. For a phage like lambda, where transcriptional organization has been unambiguously established by rigorous genetics and molecular approaches (though new technologies are providing new information ^72^), these arguments may not hold. The simplest possibility is that there are large variations in the effectiveness of each roadblock^73^, despite the perfect match of 28 nucleotides in each crRNA and, in each case, a TTTV PAM sequence. Hence, in the absence of data assessing the level of read-through in the *orf64* and *R* roadblocks, useful interpretation of the pR’ results is not practical. An intriguing possibility is that Q-mediated anti-termination may play a role in read through of these CRISPRi roadblocks. It is widely unappreciated that for all the well-studied phages, late gene expression is always under positive control, either by an antiterminator like lambda Q and the pR’ promoter, or by a transcription factor, like Lpa and the 11 late promoters of P1^17,47^. It would be interesting to determine quantitatively how such positive control factors affect the efficacy of Cas12a in CRISPR defense and dCas12a in roadblock knockdowns, with an eye towards possible evolutionary interactions. In any case, the results from knockdowns in the three major transcripts of lambda show that only *N, P, Q*, and *J* can be confidently established as essential genes. Thus, as noted earlier^61^, the nature of CRISPRi roadblock polarity means that essentiality can only be assigned for the last required gene on a transcript. The two major lessons from our work on lambda are; first, CRISPRi polarity could assign false positive gene essentiality and therefore recommend caution when applied to less studied phages; and second, CRISPRi based on DNA roadblocks is of limited utility for analysis of phages that, like lambda, feature long polycistronic transcriptional units. However, for the more utilitarian goal of identifying significant swaths of the phage genome that could be considered ‘non-essential’ and thus available for engineering, this approach still has high value. All of the 14kb pL transcript beyond N, comprising 15 genes, score unambiguously as non-essential.

Among phage genomes, lambda is arguably the best characterized transcriptional system because of its simplicity, with only three promoters involved in lytic development. P1 stands in stark contrast, with at least 45 transcriptional units, including 15 monocistronic units, and several genes served by both early and late promoters. In general, similar results were obtained from the genome-wide knockdown approach as with lambda (Fig. 4, Table 2). Of the 31 genes assigned essential character in the extensive P1 literature, all but 5 were detected by the knockdown screen. However, consideration of the transcript structures and gene positions reveals that of the 26 genes that read out as essential, 18 are located upstream of a gene known to be essential and thus the knockdown read-out is uninformative. Moreover, as in the case for the promoter-proximal genes in the lambda late transcriptional unit, P1 has a confounding transcript. Genes *25* and *26*, which were discovered as amber mutants and thus must be considered as known essentials, both score as non-essential genes. This constitutes a double contradiction, not only in the failure to detect essential character but also not exhibiting polarity on the cluster of genes downstream (genes *7, 24, 6*, and *5*) that correctly read-out as required cistrons. The simplest notion is that for some reason, neither the *25* or *26* roadblocks are effective. Quantitative assessment of road-block readthrough is beyond the scope of this initial validation screen, but it would be useful to determine the level of blockage and readthrough throughout the lambda and P1 libraries (as recently reported for *E. coli^73^*). This is especially true since the two confounding cases (genes *orf64* and *R* in lambda; genes *25* and *26* in P1) are at the 5’ end of a polycistronic transcriptional unit. Unlike other CRISPRi systems, the dCas12a roadblocks are reported to be independent of promoter-proximity, but that lesson has only been addressed within the *lacZYA* cistron^38^, and not for very long transcripts or for transcriptional units under the positive control of an antiterminator.

Because of tightly overlapped and transcriptionally linked genetic elements in phages, such polarity effects may be difficult to overcome using CRISPRi. The catalytically inactive version of recently reported RNA-targeting Cas13 system might solve some of the polarity effect issues associated with DNA-targeting Cas systems by modulating translation of single genes encoded within operons^29,31^. In addition, the absence of PAM requirements for Cas13 targeting and its broad-spectrum phage targeting capability may enable designing multiple gRNA targeting the same genomic locus, to quickly and comprehensively map gene essentiality landscape in diverse phages^29^. Nevertheless, in contrast to classical genetic methods such as recombineering, that require cumbersome cloning of long homology arm pairs followed by plaque screening to identify edited phages that exist at low abundance relative to wild-type, arrayed CRISPRi assay as presented here offers a simpler approach that only requires cloning a set of short gRNA sequences. By using pooled gRNAs, it may be possible to extend the CRISPRi technology to carry out pooled fitness assays and identify phage genes important in the phage life cycle in a single rapid assay.

Even though the gene essentiality mapping results are dependent on the experimental settings and conditions used in the assay systems, they do open up interesting questions and avenues to assess the role of non-essential and accessory genes in phage development and infection pathways^15,17,18^. By adopting high-throughput CRISPRi assays to map phage gene essentiality in different conditions^74^, it may be possible to study the role of such conditional gene essentiality in phage infection. Extending such studies to non-model, non-dsDNA phages may further provide us with deeper information needed to study genomic architecture and phage engineering applications. Considering that the different CRISPR-based tools have been successfully applied to multitudes of microbial species^34,75^ and have been used to engineer diverse phages, we expect CRISPRi technology to serve as a powerful approach to rapidly identify non-essential and accessory genes and pathways in phage infection cycles.

## Acknowledgments

The authors thank Dr. Jennifer Doudna (Innovative Genomics Institute, UC Berkeley) for sharing reagents and guidance. The authors also thank Dr. Benjamin Adler (Innovative Genomics Institute, UC Berkeley) for comments on the early draft of this manuscript. This work was funded by ENIGMA, a Scientific Focus Area Program supported by the U.S. Department of Energy, Office of Science, Office of Biological and Environmental Research, Genomics: GTL Foundational Science through contract DE-AC02-05CH11231 between Lawrence Berkeley National Laboratory and the U.S. Department of Energy. Phage P1 assay part of this research was supported by the DOE Office of Science through the National Virtual Biotechnology Laboratory, a consortium of DOE national laboratories focused on response to COVID-19, with funding provided by the Coronavirus CARES Act and the early design part of this project was funded by the Microbiology Program of the Innovative Genomics Institute, Berkeley (to A.P.A, and V.K.M.). R.F.Y acknowledges funding from NIGMS grant R35GM136396.

## Author contributions

D.P., A.P.A. and V.K.M. conceived the project. D.P. designed and built the CRISPRi libraries. D.P. N.N., M.L.M., L.R.H. and V.K.M. performed experiments. D.P., A.P.A. and V.K.M. analyzed the data. B.F.C and R. Y. provided critical reagents, insights and guidance. D.P., R.Y., A.P.A. and V.K.M. wrote the paper. All authors read and edited the paper. V.K.M., and A.P.A. arranged funds and supervised the project.

## Competing Interests

V.K.M. is a co-founder of Felix Biotechnology. A.M.D. is an advisor to Felix Biotechnology. A.P.A. is a co-founder of Boost Biomes and Felix Biotechnology. A.P.A. is a shareholder in and advisor to Nutcracker Therapeutics. The remaining authors declare no competing interests.

## Methods and materials

### Bacterial strains and phages

The bacterial strains and phages used in this study are listed in Supplementary Table 1. The oligonucleotides used in this study are listed in Supplementary Table 2. All enzymes were obtained from New England Biolabs (NEB) and oligonucleotides were received from Integrated DNA Technologies (IDT). Unless noted, all strains were grown in LB supplemented with appropriate antibiotics at 37 °C in the Multitron shaker. All bacterial strains were stored at −80 °C for long-term storage in 15% sterile glycerol (Sigma). The genotype of *E*. *coli* strains used in the assays include BW25113 (K-12 *lacI+rrnBT*14 Δ(*araB–D*)567 Δ(*rhaD–B*)568 Δ*lacZ4787*(::*rrnB*-3) *hsdR*514 *rph*-1).

*E. coli* strains were cultured in LB (Lennox) [10 g/L Tryptone, 5 g/L NaCl, 5 g/L yeast extract] or LB agar [LB (Lennox) with 1.5% Bacto agar)] at 37°C. *E. coli* strains transformed with plasmids were selected in the presence of 100 ug/mL ampicillin (LB Amp) or 30 ug/mL chloramphenicol (LB cam). Phages were plated using 0.5% top agar [10 g/L Tryptone, 10 g/L NaCl, 5 g/L Bacto agar]. 5 mM CaCl2 and 5 mM MgSO4 were added to top agar aliquots before plating.

The phages (lytic phages, λcI857 and P1vir) used in this study were prepared by the confluent plate lysis method using LB bottom plates and 0.5% top agar^76^. Phages were harvested in SM buffer (Teknova), filter-sterilized and stored at 4°C. Plaque assays were performed using spot titration method^76^

### Design and construction of spacer duplex

Cas12a recognizes TTTV as the PAM site^48^. For each target gene, PAM sites for Cas12a were identified to serve as toe-holds for the crRNAs. As any genes could have an alternative start site, the PAM sites nearby the annotated start codon of the gene were avoided. To avoid end effects, and based on prior experience in bacterial CRISPRi^65^, PAM sites were prioritized if they occurred after 20% of the gene length (so that the dCas12a complex would bind to approximately on the 1/5th position of the gene). The 28 bp nucleotide sequences immediately downstream of the PAM site in the coding strand were selected as the protospacer region. The forward oligo was designed by adding sequences “AGAT” to the 5’ region of the protospacer sequence and sequence “G” to the 3’ region of the protospacer sequence to make the ends of oligos Golden Gate cloning compatible. The reverse oligo was designed by reverse complementing the protospacer sequence from the coding strand and adding sequences “GAAAC” to the 5’ end. Custom python scripts (https://github.com/NickNolan/phage-crispri) were designed for identifying the protospacer regions and respective oligonucleotides.

We processed oligonucleotides by carrying out 5’ phosphorylation and annealing of complementary oligonucleotides in a single tube reaction. The published sequences for phages P1 (NCBI Reference Sequence: NC_005856.1) and λ (NCBI Reference Sequence: NC_001416.1) were used as reference sequences to generate oligos. Each 5 uL reaction comprised 0.5 uL each of the forward and reverse oligonucleotide pair (100 uM stock), 0.5 uL of 10x T4 DNA Ligase Reaction Buffer (NEB), 0.5 uL T4 Polynucleotide Kinase (NEB). The reaction was carried out in a thermocycler as follows: 37°C for 30 mins, 95°C for 5 mins, followed by gradient decrease of temperature from 95°C to 25°C (0.5°C every six seconds for 140 cycles). To make a working stock of the spacer duplex, the reaction mix was diluted to a final volume of 100 uL by adding milliQ water.

### Plasmid construction

The plasmid collection used in this study is listed in Supplementary Table 3. All plasmid manipulations were performed using standard molecular biology techniques. The plasmid system encoding nuclease active LbCas12a has been described previously^48^. In brief, LbCas12a is cloned under anhydrotetracycline (aTc)-inducible Tet promoter, whereas the CRISPR arrays are constitutively transcribed from a strong, synthetic promoter proD^48^. For CRISPRi, catalytically deactivated LbCas12a (dLbCas12a) lacking endonuclease activity was generated by the mutating nuclease domain of LbCas12a. For each CRISPRi plasmid, a spacer targeting a specific phage gene was cloned into the CRISPR array using Golden Gate assembly^77^. Each 5 uL of the reaction contained 0.5 uL of ATP (NEB), 0.5 uL DTT (1 mM final concentration), 0.5 uL 10x CutSmart Buffer (NEB), 0.375 uL BbsI (NEB), 0.125 uL T4 Ligase (NEB), 20 fmol CRISPRi plasmid and 100 fmol spacer duplex (0.2 uL of the working stock of the spacer duplex). The reaction was cycled between 37°C and 20°C for 5 mins each at each temperature for 30 cycles and heat inactivated at 80°C for 20 mins. This same method was followed to clone the spacer duplex into the plasmid encoding nuclease active version of LbCas12a.

For inserting a random DNA barcode into a non-essential region of phage, a recombination template was constructed on pBAD24 vector backbone^78^. A synthetic dsDNA was obtained from IDT as a gBlock gene fragment that comprised two homology arms, each of 100 bp homology to the non-essential region of the phage genome^79^. In between the two 100 bp homology arms, a random 20bp DNA barcode flanked by two primer binding regions was inserted so that the barcoded phage genome could be assayed by high-throughput DNA barcode sequencing (BarSeq) technology^69^. The gBlock fragment was PCR amplified and cloned into a PCR amplified pBAD24 backbone using Gibson assembly^80^.

The Golden Gate or Gibson assembly mixture was transformed into competent *E. coli* 5-alpha cells (NEB) following manufacturer’s recommendations and selected by plating on LB in the presence of appropriate antibiotics. Successful insertion into the plasmid backbone was verified by sanger sequencing (UC Berkeley DNA Sequencing Facility or Elim Biopharmaceuticals, Inc.). These pBAD24 derived plasmids would serve as recombination templates.

### CRISPRi assays for mapping phage gene essentiality

For CRISPRi knockdown assays, each variant of the CRISPRi plasmid was transformed into *E. coli* str. BW25113 using standard method^81^ and selected on independent LB cam plates. An overnight culture of the transformed strain was used to prepare a lawn on LB cam supplemented with 2nM or 4nM aTc for induction of the dCas12a. Phages were serially diluted ten-fold and 2 uL of each dilution was plated on a lawn of bacterial host. The number of plaques was quantified after overnight incubation at 37°C. The efficiency of plating (EOP) was calculated as the ratio of plaques appearing on BW25113 lawn expressing gRNA targeting respective phage genes to plaques appearing on BW25113 lawn expressing non-targeting gRNA. The complete compendium of EOP for each CRISPRi knockdown assay for lambda and phage P1 is listed in Table 1 and 2 respectively.

To assess the essentiality of the Nin region in λcI857, we transformed all *nin* targeting CRISPRi plasmids into *E. coli* str. BW25113, carrying a pQ plasmid system^67^ and carried out CRISPRi knockdown assays as described above (Supplementary Fig. 2). The plasmid pQ, a low-copy plasmid carrying Q, which encodes the λ late gene activator under control of a *lac*/*ara* hybrid promoter, which is inducible with IPTG and arabinose.

### Engineering DNA barcoded phages

For inserting the DNA barcode into the phage genome, pBAD24 derived plasmid was transformed into *E. coli* str. BW25113 using one-step transformation method^81^. Phage stock was appropriately diluted and plated on the lawn of the transformed BW25113 host using full-plate titration method^76^. Individual plaques were picked from the lawn and the insertion of the DNA barcode was verified by PCR amplifying the junction and sanger sequencing. The phages obtained from each plaque had a mixed population of unmodified and recombinant phage. This mixed population of phages were further enriched by confluent lysis plating method and the wt phage in each plaque was counter-selected by plating the mixture phage on the lawn of BW25113 host expressing nuclease active Cas12a target the non-essential region of the phage^82^.

### Barseq assays using DNA barcoded phages

To demonstrate the utility of barcoded phages, we mixed uniquely barcoded P1 and Lambda phage lysates in different ratios, in triplicates. To perform Barseq PCR reactions, we used phage lysates as templates. BarSeq PCR in a 50 µl total volume consisted of 20 µmol of each primer. We used an equimolar mixture of BarSeq_P2 primers along with new Barseq3_P1 primers as detailed earlier^69,83^. Briefly, the BarSeq_P2 primer contains the tag that is used for demultiplexing by Illumina software, and the new Barseq3_P1 primer contains an additional sequence to verify that it came from the expected sample (as described earlier)^83^. All experiments done on the same day and sequenced on the same lane. Equal volumes (5 µl) of the individual BarSeq PCRs were pooled, and 50 µl of the pooled PCR product was purified with the DNA Clean and Concentrator kit (Zymo Research). The final BarSeq library was eluted in 40 µl water. The BarSeq libraries were sequenced on Illumina HiSeq4000 instrument with 50 SE runs. We used in-house Barseq PCR processing code for estimating DNA barcodes in samples^69^.

### Data availability

Supplementary Tables 1-3 are deposited here: https://doi.org/10.6084/m9.figshare.22817084.v1

---

**Supplementary Figure 1.**
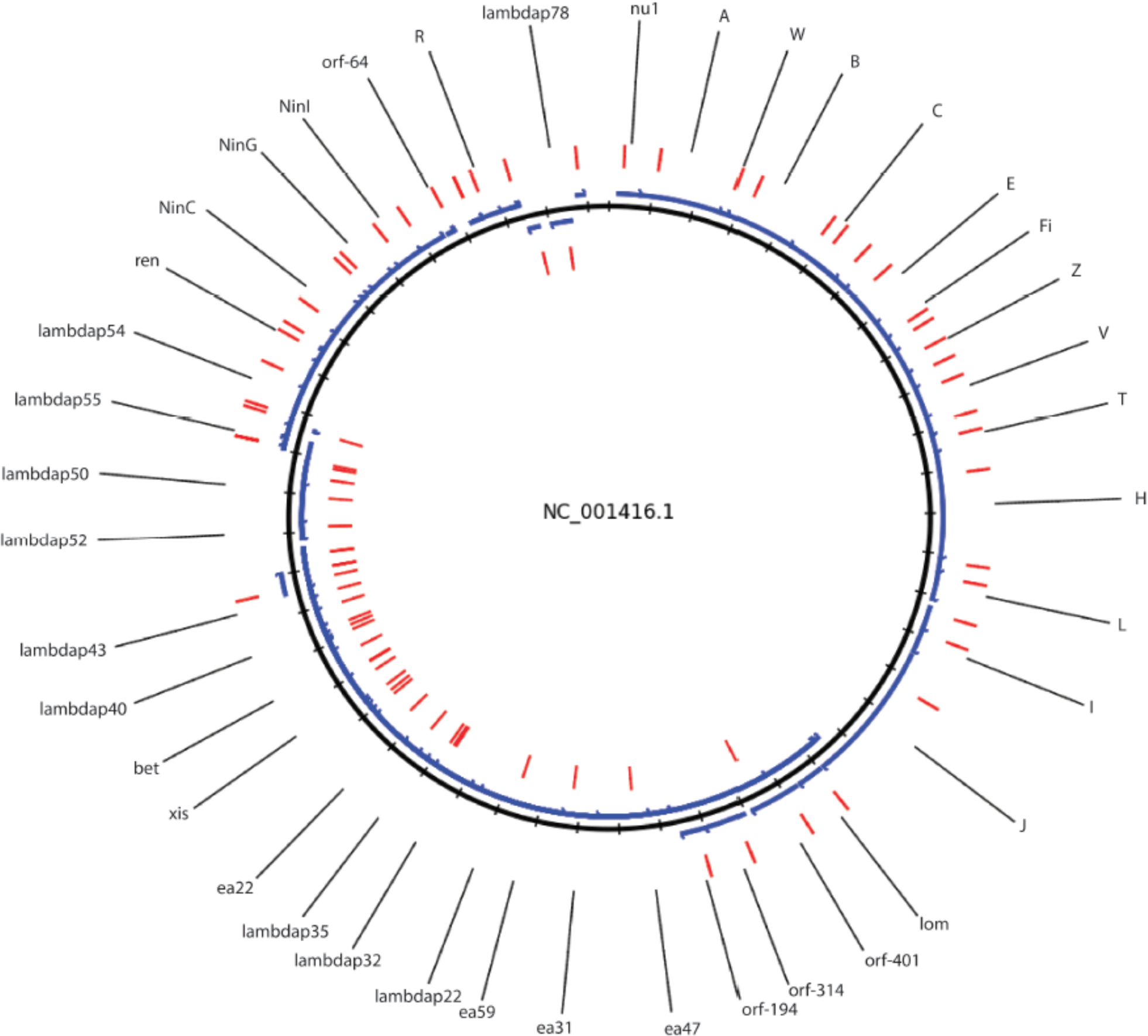
**Lambda phage genome**, CRISPRi oligo designs. blue represents genes, red represents primers, and black is the full genome. Outside is on the positive strand, where inside is negative

**Supplementary Figure 2.**
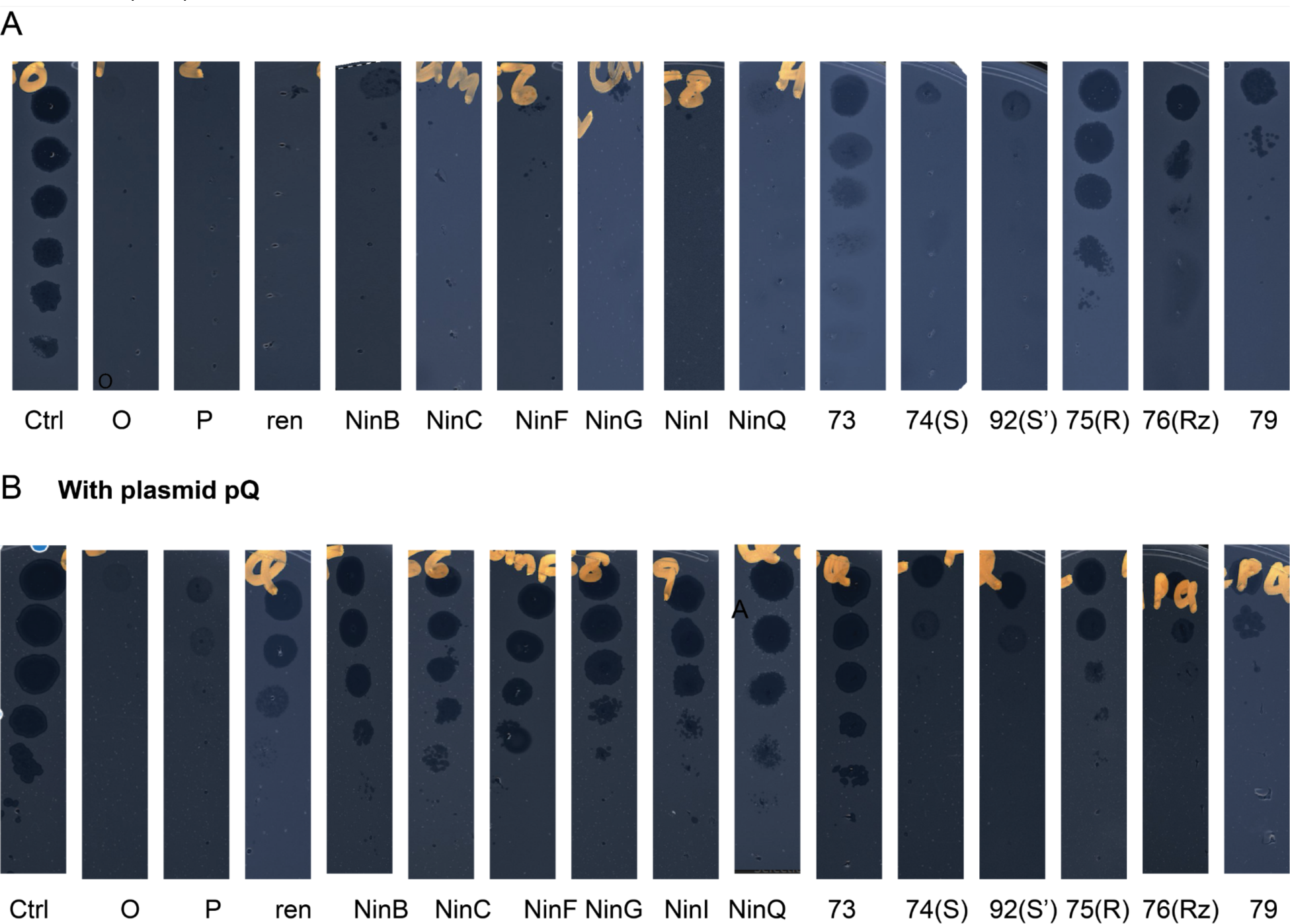
**CRISPRi of Nin region without and with plasmid pQ expressing gene Q:** EOP experiments for assessing CRISPRi polarity effect on Nin region. (A) EOP for CRISPRi assay for each gene shown (each strain with individual CRISPRi plasmid. (methods). (B) EOP experiments in presence of plasmids pQ and CRISPRi targeting each gene in Nin region context. Assays were performed in BW25113 strain expressing a non-targeting crRNA as a control (Ctrl). Inductions were done as detailed in the methods section.

**Supplementary Figure 3.**
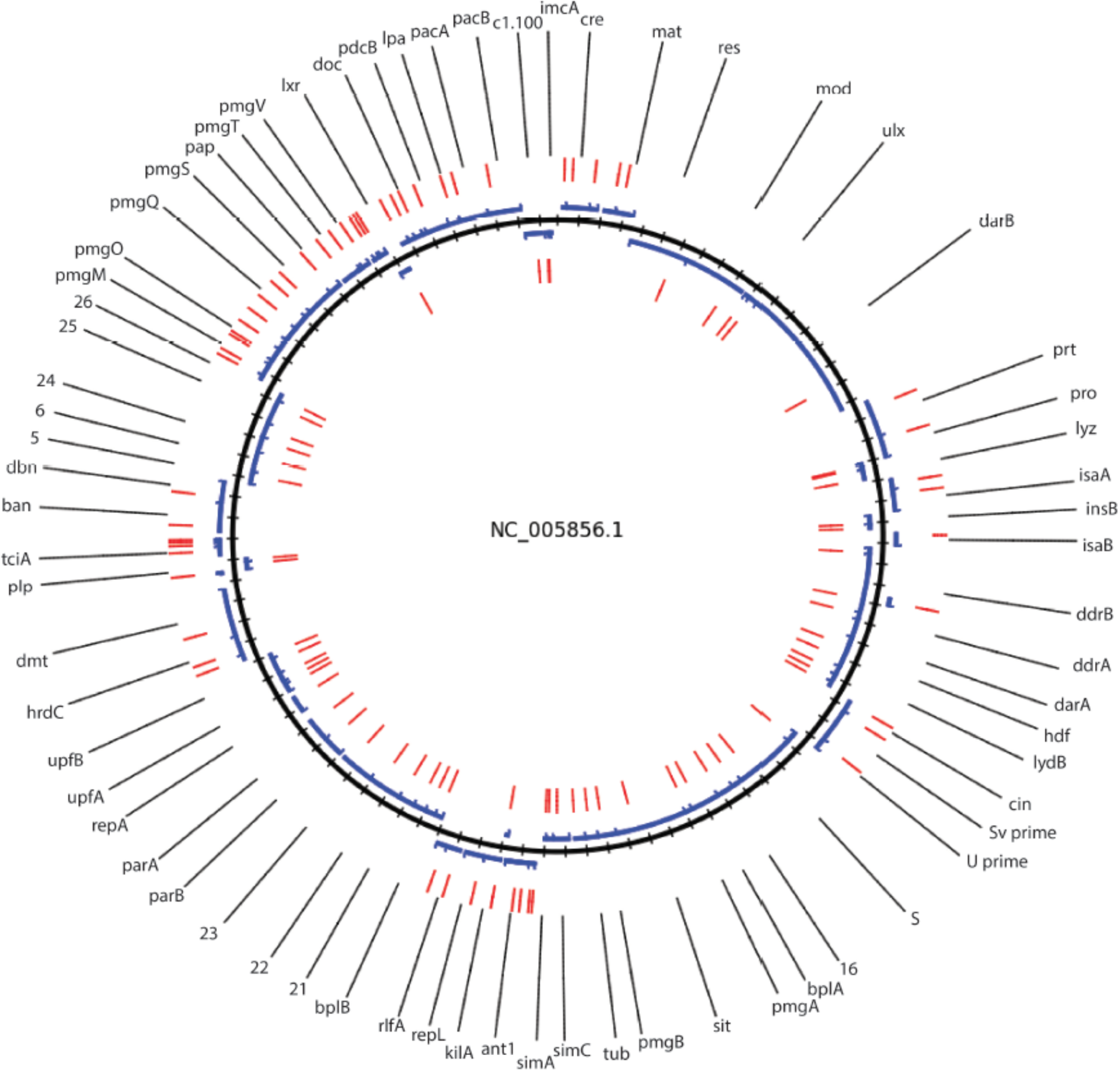
**P1 phage genome**, CRISPRi oligo designs. blue represents genes, red represents primers, and black is the full genome. Outside is on the positive strand, where inside is negative

## Notes

https://doi.org/10.6084/m9.figshare.22817084.v1

### Summary of Updates

Corrected author list order

